# Herpes ICP8 protein stimulates homologous recombination in human cells

**DOI:** 10.1101/259739

**Authors:** Melvys Valledor, Richard S. Myers, Paul C. Schiller

## Abstract

Recombineering has transformed functional genomic analysis. Genome modification by recombineering using the phage lambda Red SynExo homologous recombination proteins Beta in *Escherichia coli* has approached 100% efficiency. While highly efficient in *E. coli*, recombineering using the Red SynExo in other organisms declines in efficiency roughly correlating with phylogenetic distance from *E. coli*. SynExo recombinases are common to double-stranded DNA viruses infecting a variety of organisms, including humans. Human Herpes virus Type 1 (HHV1) encodes a SynExo comprised of ICP8 synaptase and UL12 exonuclease. In a previous study, the Herpes SynExo was reconstituted *in vitro* and shown to catalyze a model recombination reaction. Here we describe stimulation of gene targeting to edit a novel fluorescent protein gene in the human genome using ICP8 and compared its efficiency to that of a “humanized” version of Beta protein from phage λ. ICP8 significantly enhanced gene targeting rates in HEK 293 T cells while Beta was not only unable to catalyze recombineering but inhibited gene targeting using endogenous recombination functions, despite both synaptases being well-expressed and localized to the nucleus. This proof of concept encourages developing species-specific SynExo recombinases for genome engineering.

**SIGNIFICANCE:** Genome modification by recombineering using SynExo viral recombination proteins has transformed functional genomic analysis in bacteria. Single-stranded DNA (ssDNA) recombineering approaches 100% efficiency in *E. coli* using Beta protein from bacteriophage lambda, but recombineering has not been extended to eukaryotic genomes. Efficient recombineering requires SynExos that co-evolved with a viral host, however SynExos are common to viruses infecting a variety of organisms, including humans. The ICP8 protein of Human Herpes virus Type 1 is a SynExo protein similar to Beta. In this pioneering study, Herpes ICP8 stimulated gene targeting in a human genome by homologous recombination while the bacterial virus Beta protein inhibited recombination in human cells. This is the first demonstration of host-specific recombineering in human cells using a human viral SynExo protein.

## INTRODUCTION

Bacterial recombineering catalyzes highly efficient homologous recombination using viral two-component recombination modules we named SynExo complexes (Vellani & Myers 2003). The exonuclease component of the SynExo resects linear dsDNA substrates to create single-stranded DNA (ssDNA) ends. The synaptase component of the SynExo is loaded onto the nascent ssDNA to form a protein-DNA filament that catalyzes the search for homology and subsequent annealing to the homologous DNA target. The annealed recombination intermediate can prime DNA synthesis using the 3’ end of the invading ssDNA and the paired template DNA. Ultimately, the invading ssDNA becomes covalently joined to the targeted DNA and stably transmitted as a permanent part of the target DNA. Recombineering has become instrumental in analyzing gene function in bacteria since genome modification can be seamless and is specific and highly efficient, especially when one simplifies the reaction by using ssDNA oligonucleotides (ssDNA oligos) as substrates and the synaptase component of the SynExo (Ellis et al. 2001; Zhang et al. 2003). A model for ssDNA recombineering is illustrated in Fig.1.

**Figure 1.**
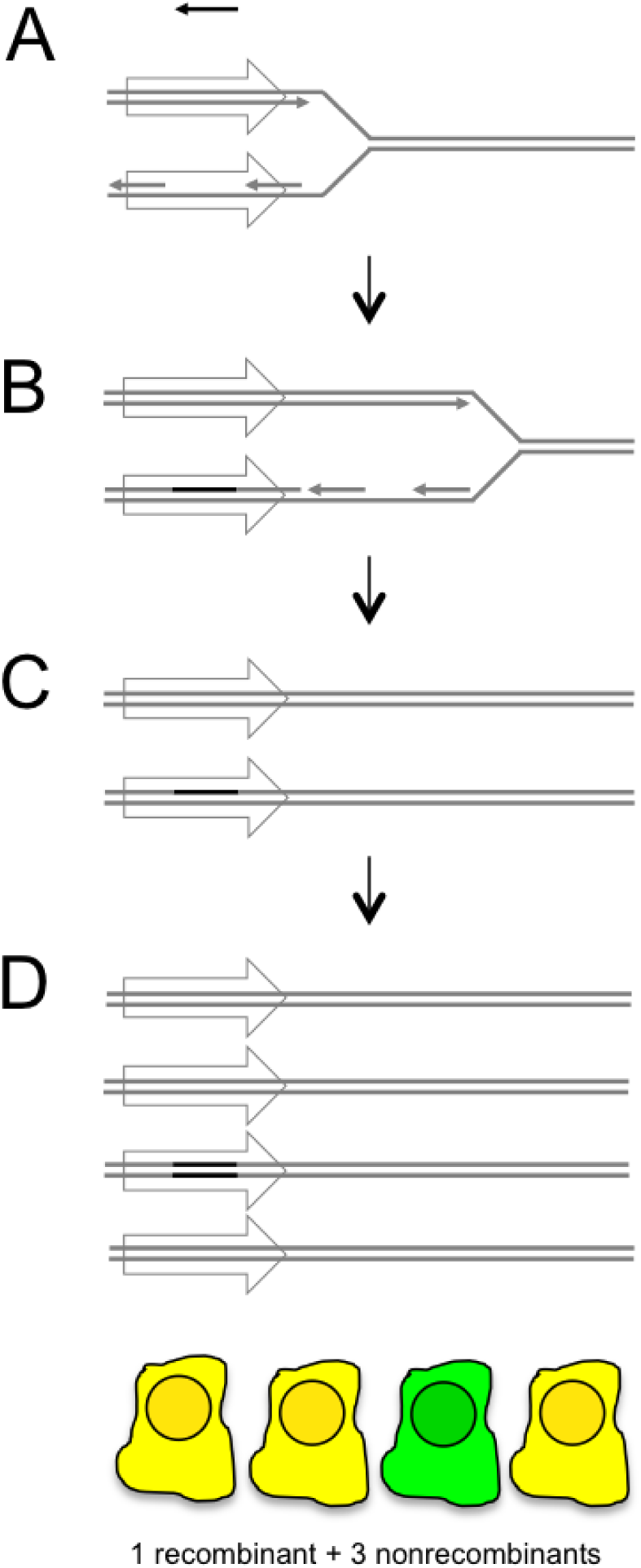
ssDNA (oligo) recombineering model. A) Synaptases bind to ssDNA (represented by the left-facing arrow at top) to form a nucleoprotein filament. B) The nucleoprotein filament samples ssDNA sequences between Okazaki fragments in replicating DNA. Synaptases promote annealing of the nucleoprotein filament to complementary ssDNA. C) The ssDNA oligo becomes joined to replicating DNA between Okazaki fragments. Completion of replication yields one nonrecombinant chromosome and one recombinant heteroduplex chromosome. D) A second round of replication and division yields one fully recombinant cell (green) and 3 nonrecombinant cells (yellow) from a cell bearing one initially recombinant target chromosome.

Eukaryotic genome segments have been recombineered with high efficiency in bacteria, but not yet in eukaryotic cells. Efforts to move enterobacteria phage SynExos into other organisms have had mixed success, depending on the phylogenetic distance from *E. coli*. For example, *Shigella* and *Salmonella* genomes have been successfully engineered using the λ Red SynExo to perform epitope tag insertion, deletions and site-directed mutagenesis (Murphy 2012), but recombineering efforts using enterobacteria phage SynExos in organisms more distantly related to *E. coli* have met less success (Datta et al. 2008; van Kessel & Hatfull 2008). Very low efficiency genomic ssDNA recombineering has also been reported in mouse embryonic stem cells using Beta protein from bacteriophage lambda (Dekker et al. 2006), although it was not clear from the reported data how well Beta was expressed and whether Beta was properly localized to the nucleus in the mouse cells. However, the negative correlation of phylogenetic distance between the host origin of the viral SynExo and the target organism with the efficiency of recombineering suggests that it may be necessary to maintain interactions between coevolved viral SynExo proteins and unknown host factors for efficient recombination.

SynExo recombinase genes appear to be common to genomes of double strand DNA (dsDNA) viruses with no RNA stage (Myers & Rudd 1998; Aravind et al. 2000). Putative SynExo components have been found in viruses that infect most types of bacteria and many eukaryotes (Fig. S1). Efforts to develop alternative SynExos (*e.g.* Table S8) that are “evolutionarily tuned” to engineer specific genomes have yielded encouraging results (van Kessel & Hatfull 2007; van Pijkeren & Britton 2012; Swingle et al. 2010). We propose that in order to extend the power of recombineering to human cells, a human viral SynExo should be employed. We previously identified a SynExo in human Herpes virus type 1 (HHV1) comprised of ICP8 synaptase + UL12 exonuclease (Myers & Rudd 1998) and subsequently showed (Reuven et al. 2003) that the HHV1 SynExo catalyzes strand exchange in an *in vitro* model of recombination using dsDNA. More recently, the HHV1 SynExo was reported to stimulate dsDNA break repair in rad52-deficient human 293 cells (Schumacher et al. 2012).

Since ssDNA recombineering is more efficient in bacteria and only requires the synaptase component of the SynExo complex, we tested ssDNA recombineering in human cells using the well characterized bacteriophage synaptase Beta and the human viral synaptase ICP8. We predicted that if recombineering were host-specific, a human viral synaptase would more efficiently promote gene targeting in human cells than would the enterobacteria phage synaptase Beta. Here we evaluated ssDNA recombineering in human cells in the presence of either the human viral synaptase ICP8 or the bacteriophage synaptase Beta.

## RESULTS

### Creation of fluorescent human recombineering reporter cell lines

For this proof of concept, we created a quantitative and sensitive reporter for genome editing using ssDNA oligos. We choose to change the emission spectrum of eGFP by targeting codon 203 (threonine, T) to code for another amino acid (tyrosine, Y) to yield a red-shifted fluorescent protein (referred to as Mostaza in (Valledor et al. 2012) and Yellow in this report) (Figs. 2A and 2B). Lentiviral vectors based on pDual-eGFP (Valledor et al. 2012) were created to transduce human 293 T cells with lentiviruses carrying eGFP or eGFP^T203Y^. Green (eGFP) and Yellow (eGFP^T203Y^) expressing cells could be distinguished and quantified by flow cytometry (Fig. 2C-E), which allowed us to evaluate a large number of cells in each trial (Fig. 2 F). The recombineering assay used oligo substrates that introduce 4 nucleotide substitutions and change the Yellow cell fluorescence to Green (Figs. 2A and S2). While the change from green to yellow and vice verse could be accomplished with a two nucleotide substitution, the addition of 2 additional silent substitutions created a bubble when the oligo anneals the template predicted to better evade DNA mismatch repair (MMR) (Fig. S2B). Incorporation of these silent mutations also allowed us to distinguish real recombinants from revertants or eGFP contamination.

**Figure 2.**
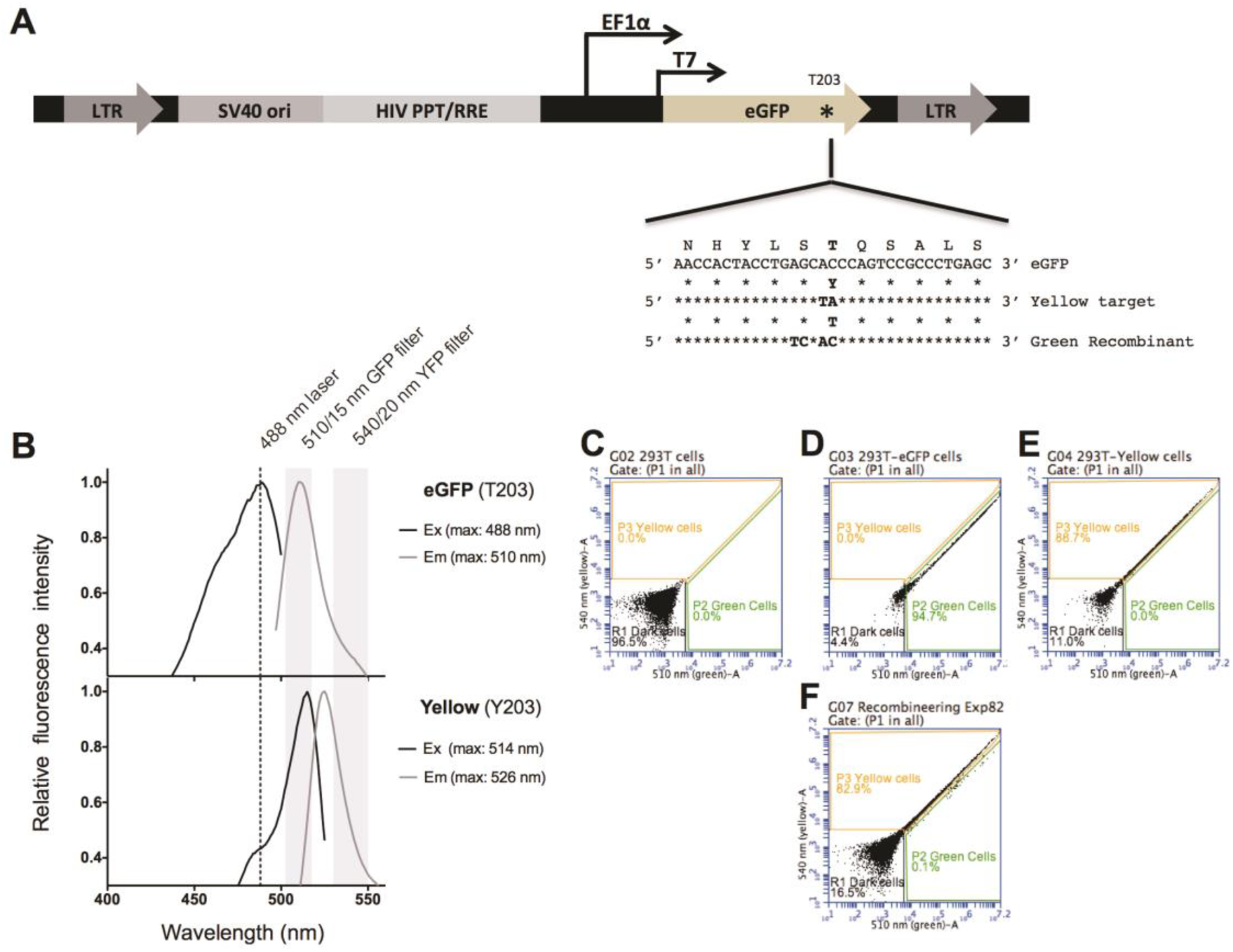
Fluorescent recombineering reporter. A) Lentiviral vector pDual-eGFP^Y203^ was used to deliver a Yellow florescent protein variant of eGFP to serve as the recombineering target transgene. B) Fluorescence excitation and emission spectra of purified eGFP and eGFP^Y203^ proteins. C-F) Flow cytometry was used to quantify 293 T-Green recombinant and 293 T-Yellow parental phenotypes. C) Untransduced cells were quantified in the R1 gate, D) pDual-eGFP transduced Green cells were quantified in the P2 gate and E) pDual-eGFP ^Y203^ transduced Yellow cells were quantified in the P3 gate. F) Example results from one recombination experiment using Yellow cells as a target and an oligo that codes for Green fluorescence.

### Sources of viral synaptases in reporter cell line 293 T-Yellow: ICP8 and HumBeta

This study aimed to evaluate if a viral synaptase can increase gene targeting rates above the endogenous rates catalyzed by host recombination proteins. While Beta protein is very efficient in bacteria, we predicted that an analogous single-strand DNA annealing protein (ICP8) from a human herpes virus would be more efficient for recombineering in human cells. Many of these studies were performed via transient transfection with ICP8 expressed constitutively from plasmid pCMV-ICP8 (used in Figs. 3, 5, 6A, S4 and S5). We noticed cell death after plasmid transfection and highly variable transfection and recombineering rates (data not shown). We subsequently created stable cell lines that expressed ICP8 and the *E. coli* bacteriophage synaptase (Beta) under the control of a doxycycline-inducible promoter. To ensure that the bacteriophage synaptase Beta would be expressed in human cells, the coding sequence was optimized for translation in mammalian cells and a nuclear localization signal was added to direct Beta to the nucleus. We called this synthetic protein “HumBeta”. In this way, we attempted to mimic an efficient bacterial recombineering protocol (Valledor 2012) where the synaptases are expressed from stable transgenes in a burst of induction just before adding the oligo substrate for recombination. By comparing recombineering efficiency using a synaptase from a virus that co-evolved with humans and a synaptase that co-evolved with a bacterium, we surmised we might probe host specificity of recombineering.

**Figure 3:**
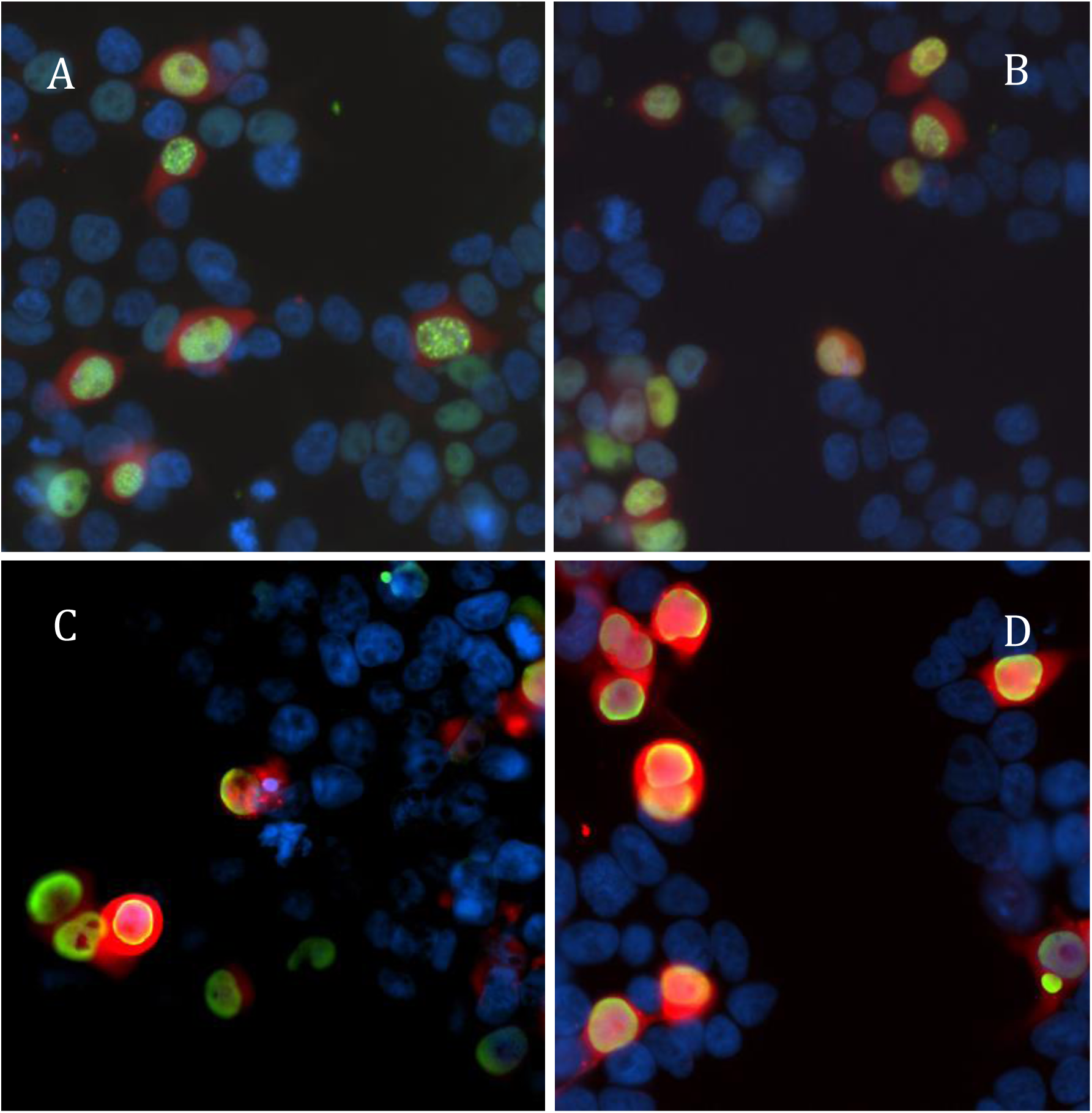
ICP8 forms punctate structures in the nucleus while HumBeta accumulates at the nuclear periphery. Transfection, induction, immunocytochemistry, and microscopy was as in Fig. S4. 293 T cells were transfected in A) with pSLIK1 (expresses PH-ICP8 with an N-terminal dipeptide), in B) with pSLIK4 (expresses PH-ICP8 with a C-terminal 18 amino acid peptide), in C) with pSLIK2 (expresses PH-NLS/HA-HumBeta with an N-terminal 41 amino acid peptide) and in D) with pSLIK5 (NLS/HA-HumBeta -V-SGS-P2A with a with an N-terminal 39 amino acid peptide and C-terminal 18 amino acid peptide). ICP8 and HumBeta are coloured green, E2-Crimson is coloured Red and DAPI is coloured blue.

Synaptase genes were fused to a red fluorescent protein gene, E2-Crimson (Strack *et al.* 2009), through a P2A linker in a single open reading frame. The P2A linker causes ribosome skipping to produce equimolar amounts of the upstream and downstream protein products (Szymczak-Workman *et al.* 2012). Six pSLIK/TREPitt lentiviral plasmid constructions were created by cloning the synaptase genes either 5’ or 3’ to E2-Crimson in separate lentiviral constructs (Fig. S3; details of plasmid and cell line construction are presented in Extended Methods in the supporting information). Lentivirus particles produced from these lentiviral vectors were used to transduce 293 T-Yellow cells. Synaptase expression was inferred from E2-Crimson fluorescence (Fig. S4) and confirmed by Western Blot (Fig. S5) and immunostaining (Fig. S4). Following induction of expression by doxycycline, ICP8 and HumBeta localized to the nucleus (Fig. S4). ICP8 was either evenly distributed or formed bright punctuate dots (Fig. 3), similar to those described in a previous report that showed ICP8 to be co-localized with pre-replication centers (de Bruyn Kops & Knipe 1988). Unlike ICP8, HumBeta did not organize into punctuate subnuclear dots, accumulating around the nuclear periphery instead (Fig. 3).

### HHV1 ICP8 protein stimulates gene targeting in human cells

Gene targeting was evaluated in 293 T-Yellow cells expressing ICP8 either from transiently transfected pCMV-ICP8 or from stably transduced pSLIK 1 & 4. Recombinant cells were identified by a shift in fluorescence from Yellow to Green using flow cytometry (Fig. 2 F). The average frequency of Green fluorescent cells among total fluorescent cells after subtracting the background from the no-oligo controls in 8 independent experiments was close to 0.2% for multiple independent ICP8 cell lines while endogenous rates were below 0.1% (data from Figs. 4–7 and Fig. S6). In Fig. 4B, human Herpes ICP8 stimulates gene targeting over endogenous recombination functions by 2.3-fold. We likely underestimated the amount of recombination promoted by ICP8 in these experiments, as it became evident that not all the cells responded to doxycycline induction. Since Crimson is expressed in equimolar amount to the synaptase, via translational coupling through a P2A linker, recombineering efficiency was measured in the subpopulation of cells expressing Crimson (Fig. 4 C-D). If recombination is measured in ICP8-expressing cells only, then gene targeting was stimulated 4.7-fold over endogenous rates.

**Figure 4.**
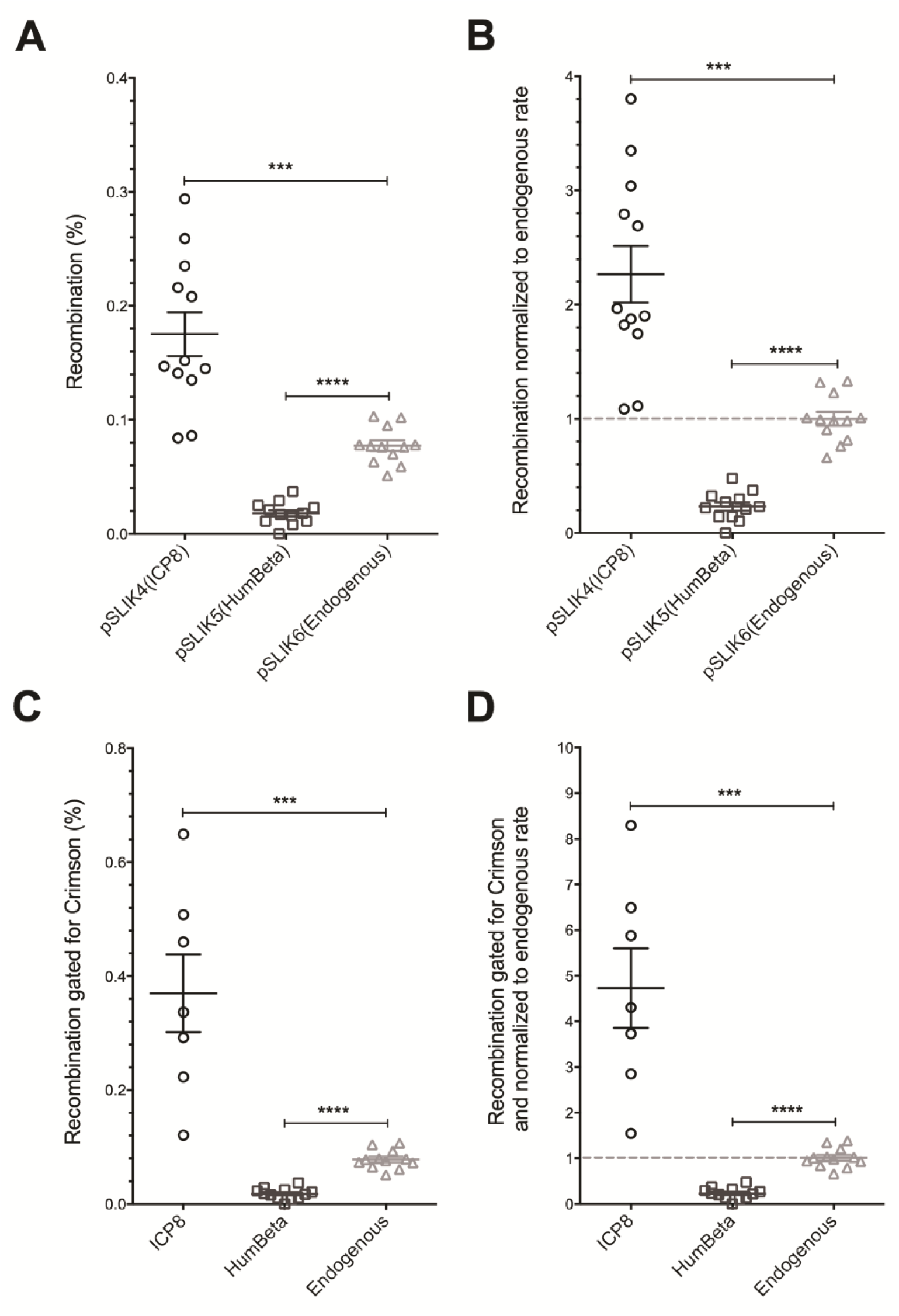
Gene targeting is stimulated by HHV1 ICP8 and inhibited by phage λ Beta in human cells. Cells were incubated with doxycycline to induce synaptase expression and subsequently transfected with 50 nM of oligo 85. A) The background of green cells in “no oligo” control experiments (0.001% for pSLIK4, 0.003% for pSLIK5 and 0.012% for pSLIK6) was subtracted from recombination frequencies. B) Recombination frequencies in (A) were normalized to the mean of the endogenous recombination frequencies in the pSLIK6 control line. C) Cells were gated for Crimson expression (which is expressed in equimolar amounts with the synaptase via translational coupling through a P2A linker) and evaluated as in (A). D) Recombination frequencies in (C) were normalized to the mean of the endogenous recombination frequencies in the pSLIK6 control line and evaluated as in (A). Data were evaluated by Fisher’s Exact Test of significance; *** indicates P<0.005 and **** indicates P<0.001. In this figure, n = 12 and error bars reflect SEM.

### Humanized Beta protein inhibits gene targeting in human cells

In parallel trials to those with ICP8, recombineering was evaluated in cells stably transduced with the humanized bacteriophage HumBeta synaptase. As in experiments with ICP8, HumBeta was induced by doxycycline and cells were subsequently transfected with oligos to assess recombination rates. The average frequency of Green fluorescent cells among total fluorescent cells after subtracting the background from the no-oligo controls was 0.02% for HumBeta cell lines produced by transduction with pSLIK2 or pSLIK5 (Figs. 4 and S6). HumBeta cell lines were almost 100% Crimson positive (pSLIK5 gated for Crimson vs pSLIK6, n=12, P=1), therefore recombineering efficiency gating cells for Crimson expression did not modified the initial outcome (Figs. 4 C-D). In striking contrast to ICP8, expression of HumBeta *inhibited* endogenous rates of gene targeting by 4.3-fold. Inhibition of endogenous recombination may result from sequestration of the oligo into non-productive Beta/DNA complexes, perhaps due to lack of host-specific protein interactions (see discussion).

### Confirmation of authentic gene targeting by ICP8

To determine if Green fluorescence in putative recombinants was due to inheriting the oligo sequence, gene targeting experiments were also performed with a Yellow “selfing” oligo with the same sequence as the target (Figs. S2 and 5A). The Yellow selfing oligo did not stimulate Green cell formation in either the presence (P=0.54) or absence (P=0.41) of ICP8. This is to be expected if oligo-mediated Yellow to Green conversion was due to recombination and not general mutagenesis. ICP8 expression increased the frequency of producing Green recombinants 2-fold above endogenous recombination (P=7.1x10-6 by Fisher’s Exact Test) when the Green oligo was used.

Evidence that the Green allele sequence was transferred from the recombineering oligo to the genome in cells expressing ICP8 was obtained by allele-specific PCR. Allele specific PCR was optimized using recombinant and parental templates (Fig. S7). Putative recombinant Green cells were enriched from non-recombinant Yellow cells by cell sorting. Green cells are expected to have at least one recombinant target copy, but might also have additional non-recombinant target copies due to multiple insertions of the fluorescent transgene. In addition, since ssDNA recombineering targets only one strand of replicating DNA, cells need to go through two cell divisions to segregate the recombinant from non-recombinant targets in a clone (Fig.1). We saw that the recombinant Green cells produced more products when amplified with a primer specific for the Green allele than with a primer specific to the Yellow allele. These results suggest that Green cells incorporated the Green oligo sequence via recombination.

Gene editing was confirmed by DNA sequencing of PCR products amplified from recombinant Green cells. The results shown in Fig. 5B, indicate that the sequence from the Green allele (TCCAC) is more prominent than the sequence of the Yellow allele (AGCTA) in Green cells. The silent mutations introduced to abrogate MMR were transferred along with the mutations that converted Yellow to Green, suggesting that Green cells did not arise by reversion. Taken together, it is evident that *bona fide* ICP8-dependent recombineering is modifying human genomic DNA at the target region as specified by the oligo sequence.

**Figure 5.**
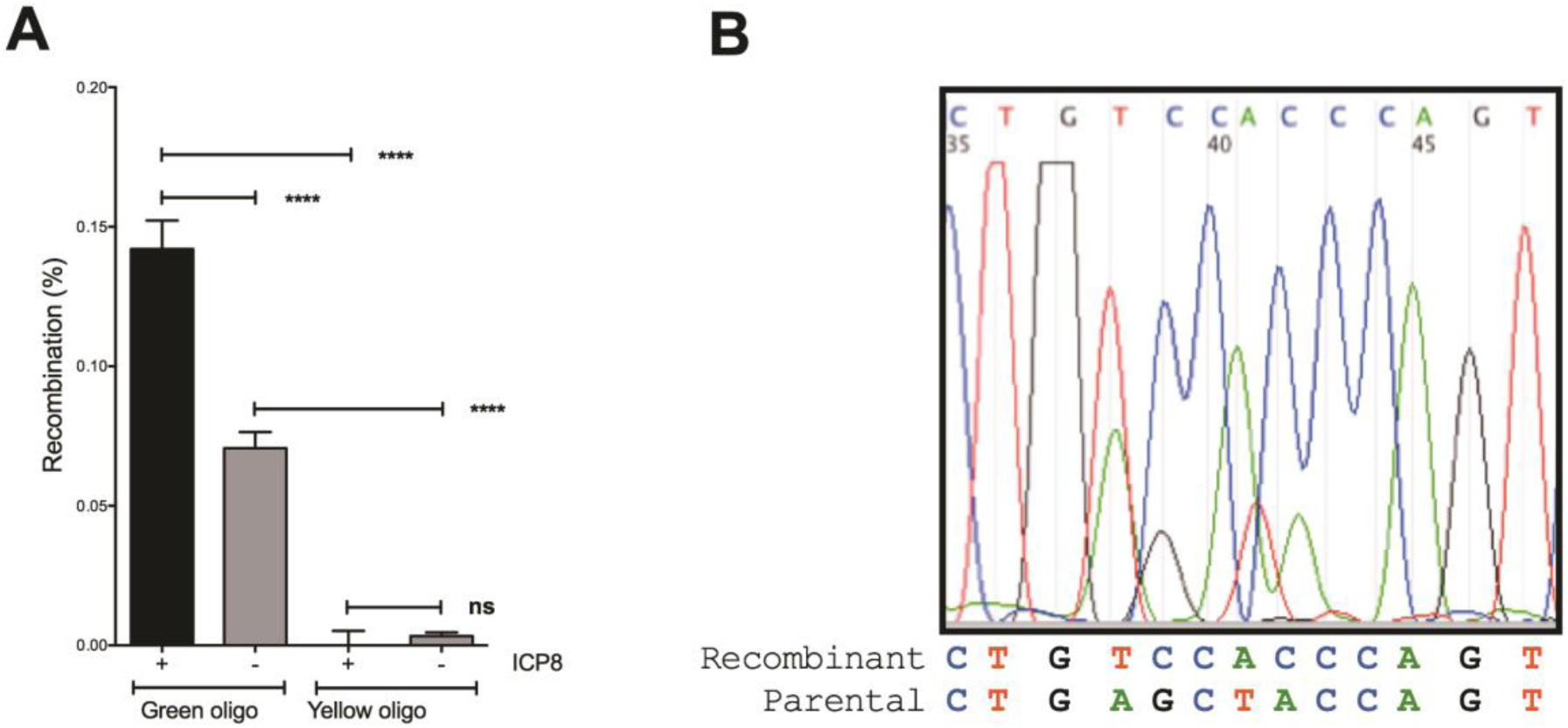
Transfer of the oligo sequence to the genome is required to change cell fluorescence. A) The Green recombinant phenotype requires a Green oligo substrate. Yellow to Green phenotypic conversion was evaluated by transient transfection with pCMV-ICP8 followed by transfection with 200 nM Green or Yellow oligos (85 and 131 respectively). B) Recombinant cells acquire the oligo sequence. A recombinant cell mixture was sorted for Green fluorescent cells. Genomic DNA was isolated from sorted cells enriched for putative recombinants. The target region was amplified by PCR and the products treated with restriction enzyme AluI to cut non-recombinant sequences. Uncut fragments were gel extracted and sequenced using primer 216. The original Parental sequence and the predicted Recombinant sequence is shown. Details of this experiment can be found in the Extended Methods under Determination of recombinant genotype.

### Oligo substrates

In bacterial recombineering, the most efficient substrates are ssDNA oligos in the range of 60 to 100 nt long (Li et al. 2003), similar to oligo lengths commonly used for gene targeting in mammalian cells (Nickerson & Colledge 2003; Andrieu-Soler et al. 2005). Shorter oligos (20–25 nt) require modification to be effective as the introduced sequences are more susceptible to exonucleases (Andrieu-Soler et al. 2005; Rios et al. 2012). To avoid potential DNA damage induced by modified DNA (Aarts & Riele 2010a), unmodified oligos of varying lengths were tested (Fig. 6A). Recombination rates improved as a function of oligo length and could be fit to a one-site binding equation with a Hill slope parameter based on previous knowledge of the cooperativity of ICP8 binding to DNA (Gourves et al. 2000). Recombination was significantly improved as oligo length increased (by Fisher’s Exact Test) in pairwise comparisons of each DNA length for a given oligo sequence in each cell line, suggesting that 65 nt oligos are better recombineering substrates in human cells than are shorter oligos. Although in our previous optimization studies in *E. coli*, little or no additional benefit was added by lengthening oligos beyond 70 nt (Valledor et al 2012), it remains to be determined if longer oligos could be more efficient for ICP8-stimulated recombineering.

**Figure 6.**
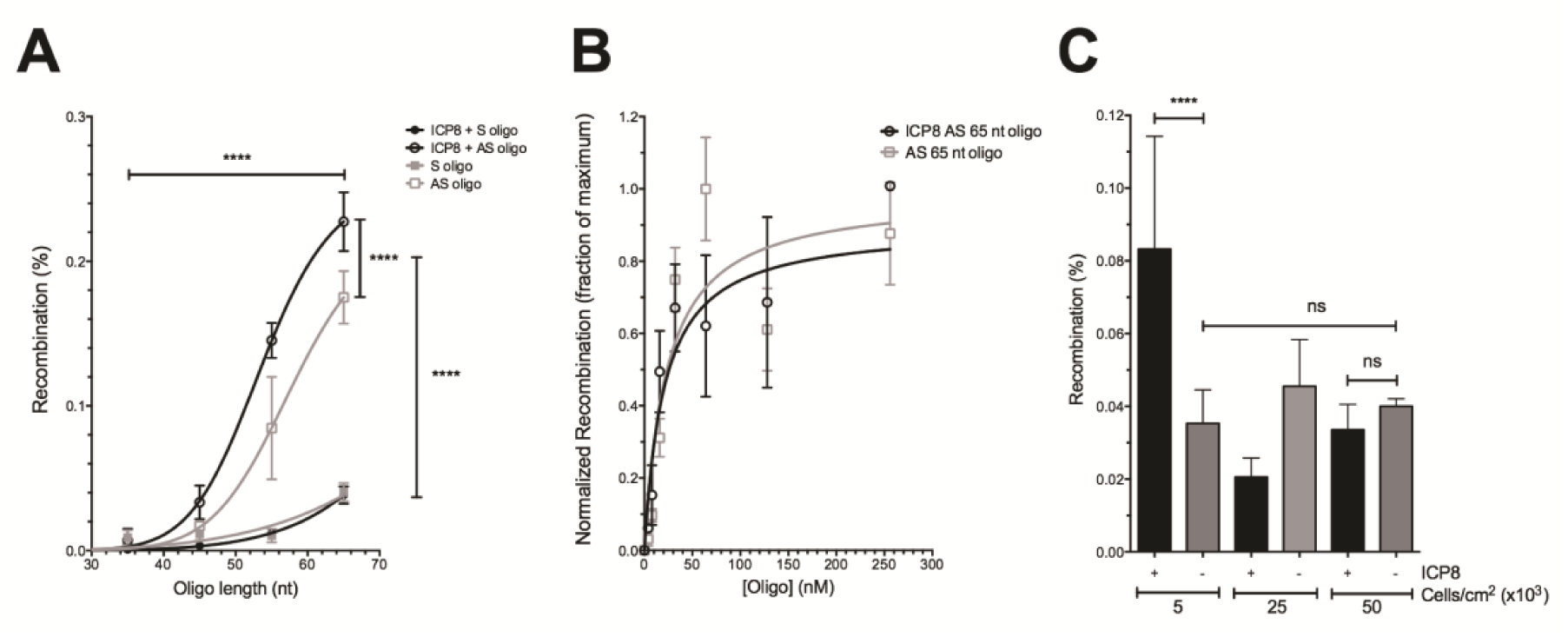
Optimization of human cell recombineering. During the course of developing this platform for human cell recombineering by ICP8, the frequency of gene targeting varied by 10 to 20 fold, influenced by the design and concentration of the oligo substrate, as well as the cell density in the well during oligo transfection. A) **ICP8 and endogenous recombination was significantly improved by longer and antisense oligos**. 293 T-Yellow cells were sequentially transfected with pCMV-ICP8 and with oligos that donate the sequence for converting Yellow to Green fluorescence. Recombination was measured as a function of oligo size and the targeted strand. AS oligo indicates that the oligo corresponds to the antisense strand of *eGFP^Y203^* and S oligo indicates that the oligo corresponds to the sense strand of *eGFP^Y203^.* The no oligo background for 293 T or 293 T-pCMV-ICP8 was subtracted from each data point and never exceeded 0.004%. Fisher’s Exact Test was used to evaluate statistical significance with **** indicating P<0.001; n=3. Recombination rates as a function of oligo length could be fit to a one site binding equation with a Hill slope parameter based on previous knowledge of the cooperativity of ICP8 binding to DNA (Gourves et al. 2000). B) **Recombination rates improved as a function of oligo concentration.** The effect of oligo concentration on gene targeting was evaluated from 4 to 256 nM. Recombination with oligo 85 was measured in pSLIK4 and pSLIK6 cell lines induced with 1 μg/ml doxycycline as a function of oligo concentration. Recombinants were quantified by flow cytometry, normalized to the maximum value for each cell line and fit to a one site binding equation that accounts for depletion of free recombination proteins at saturating oligo concentration. C) ***ICP8 requires replicating cells.*** In the course of these studies, it became evident that some of the fluctuation in recombination rates observed was due to differences in the density of cells plated before addition of oligos. Recombination with 50 nM antisense oligo 85 was measured in pSLIK4 and pSLIK6 cell lines induced with 1 μg/ml doxycycline and plated at densities of 5000, 25000 and 50000 cells per cm^2^. Recombinants were quantified by flow cytometry. Fisher’s Exact Test was used to evaluate statistical significance with **** indicating P<0.001 and ns indicating P>0.05; n=3.

Oligo uptake has been positively correlated with gene targeting frequency (Nickerson & Colledge 2003; Aarts & Riele 2010b; Andrieu-Soler et al. 2005). When the effect of oligo concentration on 293 T recombineering was evaluated by titrating a 65 nt oligo from 4 to 256 nM (Fig. 6B), recombination rates improved as a function of oligo concentration. By fitting the data to a one-site binding equation that accounts for depletion of free recombination proteins at saturating oligo, the Kd apparent for binding inferred from recombination frequency was about 2 nM, with about 96% saturation at 50 nM oligo for both endogenous and ICP8-enhanced recombination.

### ICP8 requires replicating cells

In the course of these studies, it became evident that some of the fluctuation in recombination rates observed was due to differences in the density of cells plated before addition of oligos. When cell plating density was varied from 5,000 to 50,000 cells/cm^2^, it was observed that the stimulation in recombination by ICP8 in the presence of 50 nM oligo was only seen at lower cell density (Fig. 6 C), conditions that promote more replication and consistent with the model that ssDNA recombineering targets replication intermediates (see *Strand bias* subsection, below).

### Strand bias

In bacterial recombineering and in mammalian gene targeting the most efficient substrates are oligos that target the lagging strand template (Aarts & Riele 2010a; Li et al. 2003; Valledor et al. 2012). Since the eGFP^Y203^ transgene was delivered to cells by lentiviral transduction, the reporter target gene was likely integrated in random orientations in individual cells with respect to chromosomally initiated replication forks. Therefore, we predicted that oligos targeting each strand of the Yellow target would produce recombinants at equal frequency when using pools of cells with randomly integrated target genes. Surprisingly, when either DNA strand was targeted in parallel experiments with oligos resembling either the sense or the antisense sequence of the Yellow target, the oligos with the antisense sequence consistently produced more Green cells (Fig. 6A). By subtracting the background of endogenous recombination, it became evident that only antisense oligos acted as ICP8-dependent recombineering substrates.

The observed strand bias was not likely due to MMR acting on recombination intermediates differently as the 4 nucleotide mismatch in heteroduplex recombination intermediates formed when hybridized to the lagging strand template (Fig. S2B) are predicted to evade MMR (Valledor et al. 2012) and MMR activity is largely absent in 293 T cells (Schumacher et al. 2012; Trojan et al. 2002). Likewise, the direction of transcription was not likely relevant based on previous studies of strand bias in gene targeting (Aarts & Riele 2010a; Ellis et al. 2001). It was previously seen that replication was essential for gene targeting (Olsen et al. 2005; Poteete 2013) and that the lagging strand template in DNA synthesis provided the best target for oligo-mediated recombination (Li et al. 2003; Ellis et al. 2001; Valledor et al. 2012) (Fig. 1). Since the Yellow reporter target gene was likely integrated in random orientations in each individual cell with respect to chromosomally initiated replication forks, we cannot account for this bias based on proximity and orientation with respect to endogenous origins of replication. However, the lentiviral vector pDual-Yellow contains an SV40 origin of replication located 5’ to the *eGFP*^Y203^ gene (Fig. 2A). If this SV40 origin initiated replication with the help of the T antigen expressed in 293 T cells, then we expected antisense oligos to hybridize to the lagging strand template and produce the observed recombinants. 293 T was created using a temperature sensitive variant of the SV40 T antigen that fails to initiate replication from the SV40 origin at elevated temperature. Therefore, gene targeting was examined in cell lines expressing ICP8 or only using endogenous recombination functions at 37°C (permissive for T antigen activity) and 39.5°C (nonpermissive for T antigen activity). At 37°C, the strand bias in gene targeting was evident in both the presence and absence of ICP8 (Fig. 7). However, as recombination rates were reduced and strand bias eliminated at 39.5°C, we conclude that SV40 origin-dependent replication across the target locus improved gene targeting frequencies and was responsible for the target strand bias.

**Figure 7.**
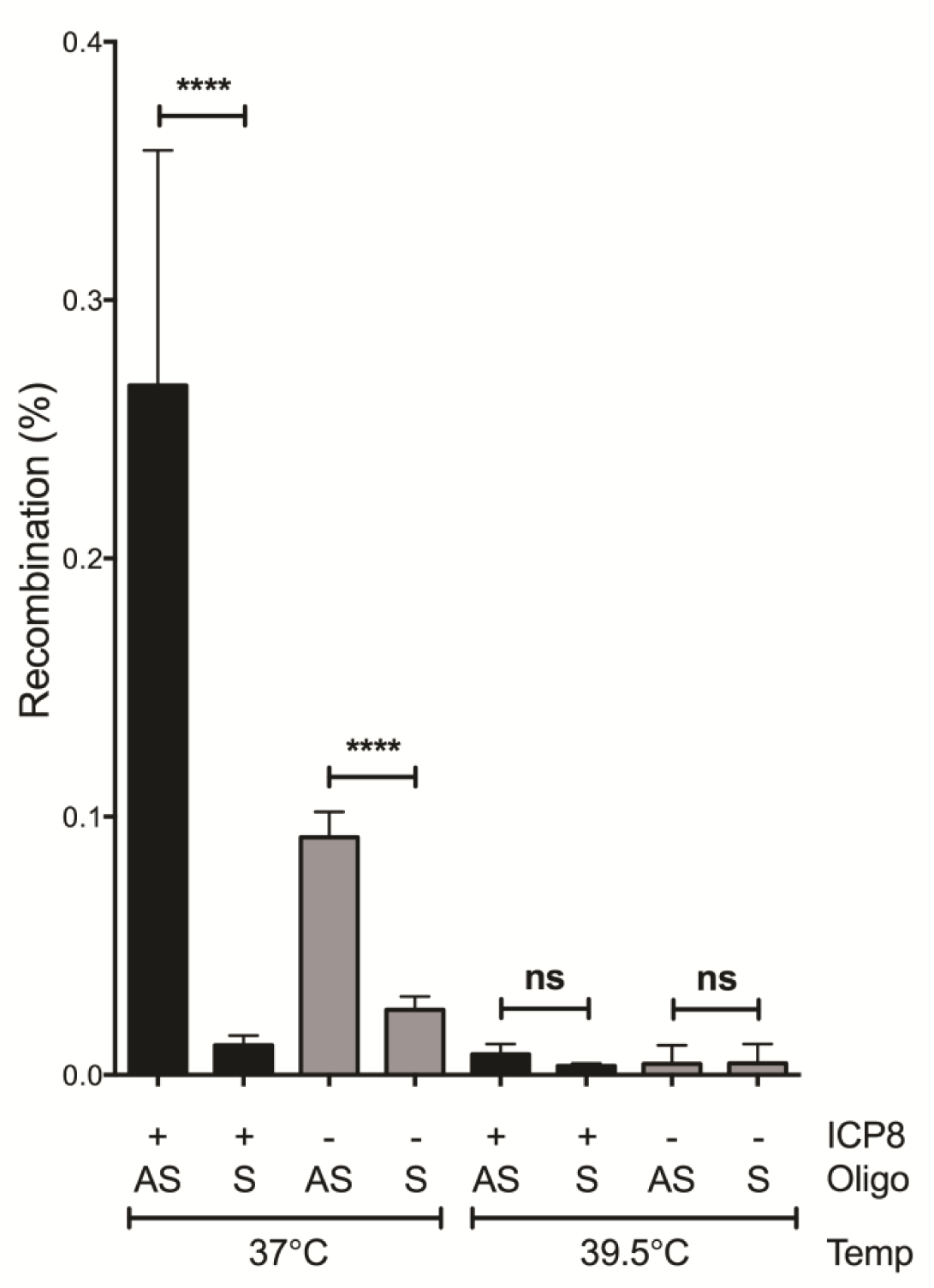
Recombineering strand bias depends on replication originating from the SV40 origin in the integrated lentiviral transgenic target. Recombination was measured in pSLIK4 and pSLIK6 lines using antisense or sense oligos. Cells were grown either at 37°C (T antigen active), or at 39.5°C (T antigen does not bind to or promote replication from the SV40 origin). Fisher’s Exact Test was used to evaluate statistical significance with **** indicating P<0.001 and ns indicating P>0.05. n=3.

## DISCUSSION

Our observations are consistent with the proposal that ICP8 is a SynExo synaptase subunit like Beta protein from bacteriophage λ. It has been previously demonstrated that Beta protein directs ssDNA recombination substrates to complementary sequences exposed at the replication fork. Our data suggest that T-antigen-dependent replication across the target gene stimulates ssDNA recombination by exposing ssDNA primarily on the lagging strand template between Okazaki fragments. Single-strand DNA gaps on the lagging strand template expose complementary sequences to the “antisense” oligos we found to be most effective at gene targeting. These results bear striking similarity to bacteriophage SynExo-mediated recombination operating in bacteria. Supporting this conclusion, we found that ICP8 promoted gene targeting only when the cells were seeded at low density and had the opportunity to replicate several times after oligo transfection. The DNA sequence of the recombinant Green fluorescent cells confirmed that genetic information in the oligo was precisely transferred to the target genome by recombineering. We predict that the acquisition of the oligo sequence reflects oligo annealing by ICP8 to the homologous target region accompanied by limited DNA synthesis and ligation to covalently join the introduced oligo to the genome. The details of this mechanism remain to be determined, but as modelled in Fig. 1, the oligo sequence once incorporated, could be either repaired by MMR or stably transmitted after a round of DNA synthesis.

Strikingly, the well-studied and commonly used bacteriophage λ synaptase Beta not only failed to stimulate recombination, but significantly inhibited gene targeting by endogenous functions. These results recapitulate recent studies where Beta expression in HT1080 cells did not promote chromosomal gene correction (Xu et al. 2015). Instead, the authors demonstrated a modest stimulation of gene editing by Beta using a co-transfected plasmid as the target replicon. When ICP8 accumulates in the nucleus, it localizes to subnuclear domains called pre-replicative sites (Taylor et al. 2003) and physically associates with several human proteins involved in DNA metabolism and chromatin remodelling (Taylor & Knipe 2004). In contrast, when “humanized” Beta accumulates in the nucleus, it fails to form the subnuclear punctate staining domains seen in ICP8 expressing cells and localizes predominantly to the nuclear periphery (Fig. 3). Images of Beta expression in HT1080 cells also show an enrichment of Beta protein near the nuclear periphery and not in the subnuclear domains called genome replication centers (Xu et al. 2015). Perhaps the failure to organize into subnuclear assemblies reflects a lack of host-specific protein interactions required to stimulate recombination and instead sequesters ssDNA substrates from endogenous recombination functions. Given that the degree of stimulation of gene editing by ICP8 in the human genome exceeds that of Beta but is similar to Beta acting on synthetic extrachromosomal DNA targets, we propose that ssDNA annealing proteins have near universal function but must be tuned to the molecular context, *e.g*. genomic chromatin.

These results are consistent with the model that viral synaptase-mediated recombineering is host-specific (Murphy 2012; Thomason et al. 2005; van Kessel et al. 2008) due to co-evolution of viral SynExo recombinases and host cell proteins. According to this model, recombineering is host-specific because viral SynExo complexes cooperate with cellular functions to promote recombineering and viral development. We suggest that SynExo interactions with host proteins are required to coordinate recombineering in replication forks. In order to achieve the high rates of recombineering obtained in finely tuned bacterial systems, the human viral SynExo recombination mechanism will likely benefit from coordination with the cell cycle.

We observed an average of ~0.1% oligo-dependent gene targeting in the absence of ICP8, likely facilitated by endogenous cellular recombination functions such as Rad52, a synaptase protein with structure and function similar to viral synaptases (Shinohara et al. 1998; Lopes et al. 2010). Both endogenous and ICP8-stimulated gene targeting in 293 T cells may have benefitted from impaired MMR (Trojan et al. 2002; Cannavo et al. 2007) and increased replication in the presence of the T antigen (Friedrich et al. 1992; Perry & Lehman 1998).

While previous data suggest that ICP8 and Rad52 partially overlap in function (Schumacher et al. 2012), we found that the human viral synaptase ICP8 significantly (albeit modestly) and reproducibly increased gene targeting rates. Although ICP8 stimulated recombination, we suspect that our results underestimate these rates. ICP8 expression was observed to reduce cell viability (data not shown), likely leading to reduced recovery of recombinants. Indeed, by gating for the remaining ICP8 expressing cells after outgrowth, recombinants were enriched, raising stimulation of ICP8-mediated recombination (from 2.3-fold to 4.7-fold in Fig. 4 C and 4D). Future efforts will capitalize on this enrichment method.

An alternative strategy might develop the endogenous ssDNA binding protein Rad52 as a recombineering tool. Rad52 appears to have originated from a viral synaptase gene (Ploquin et al. 2008) and functionally overlaps with ICP8 in recombinational repair of DNA breaks (Schumacher et al. 2012). Previous studies indicated that Rad52 overexpression might increase gene targeting rates (Kalvala et al. 2010; Di Primio et al. 2005; Takahashi & Dawid 2005) but that sustained high-level expression is detrimental to viability and recovery of recombinants (Yáñez & Porter 2002), consistent with our observations with constitutively expressed ICP8. We suggest that Rad52 could perform as a recombineering synaptase and is contributing to the endogenous “background” recombination we observed in the absence of ICP8.

Much remains to be learned about the human genome. Development of efficient and specific tools for genome editing is essential to dissect the role of specific alleles on human health and disease. Recombineering optimization in microbes elevated the efficiency from 6% (Ellis et al. 2001) to 15% of total cells, 40% of transfected cells, and 85% of transfected cells that are mismatch repair deficient (Valledor et al. 2012). Further improvement of eukaryotic cell recombineering might make this a worthy partner to CRISPR/Cas9 as a homologous recombination partner. The twin revolutions of stem cell therapy and genomics are providing the tools for precision medicine and a deeper understanding of human development. Extension of human cell recombineering to stem cell genome editing may lead to development of autologous stem cell genetic therapy and development of better disease models. Our results suggest that ICP8 is a *bona fide* synaptase in the SynExo family of viral two component recombinases (Reuven et al. 2003; Vellani & Myers 2003) and further encourages the development of organism-specific SynExos for genome editing in other species.

## Materials and Methods

### Plasmids

The pDual-eGFP(Y203) lentiviral plasmid was used to create the target for ssDNA oligo-mediated gene targeting, the reporter for recombineering. A scheme for it and the parental plasmid pDual-eGFP is represented in Fig. 2A, while their design and construction were described in (Valledor et al. 2012). pCMV-ICP8 (Taylor & Knipe 2003) was used for transient ICP8 transfection and as the source of ICP8 for lentiviral vectors. ICP8 and Beta synaptase genes were individually cloned into the pSLIK-Zeo lentivirus vector for high transduction efficiency, zeocin selection and doxycycline inducibility (Shin et al. 2006). The synaptase genes were fused to the E2-Crimson fluorescent protein gene (Strack et al. 2009) through a P2A linker (Szymczak-Workman et al. 2012) to follow synaptase expression (Fig. S3). Cloning details are described in Extended Methods.

### Oligonucleotides

Oligos were 35 to 65 nt long and complementary to the target region except for a two-nucleotide change specifying Yellow to Green and a silent two-nucleotide substitution at the adjacent serine codon (Figs. 2 and S2A). Unmodified ssDNA (oligos) were purchased from Sigma-Aldrich and their sequences are shown in Table S9.

### Cell lines

293 T-Yellow recombineering reporter cell lines were created by transducing 293 T cells with lentivirus produced from pDual-eGFP(Y203) (Valledor et al. 2012). Inducible Synaptase-Crimson cell lines were created from 293 T-Yellow cells transduced with pSLIK lentiviral clones 1–6. Transduction protocols are described in Extended Methods.

### Human cell gene targeting by recombineering

ssDNA recombineering was performed in exponentially growing 293 T-Yellow cells expressing a viral synaptase (ICP8 or HumBeta). In experiments in which ICP8 was expressed from a transiently transfected plasmid, 293 T-Yellow cells were first transfected using 6 μl Fugene 6 (Roche) and 2 μg pCMV-ICP8 per well in 6 well plates in triplicates. In stable 293 T-Yellow-pSLIK cell lines encoding ICP8 or HumBeta, cells were first induced with 1 μg/ml doxycycline. The next day, cells were seeded in 24-well plates in 0.5 ml 5% FBS-DMEM at 5,000 cells/cm^2^. The following day, cells were transfected by mixtures of Lipofectamine 2000 (Invitrogen) and unmodified oligos to a final Lipofectamine concentration of 0.27% and oligo concentration of 50–200 nM as indicated in the figures and text. Cells were passed to 6-well plates and collected for flow cytometry as they reached confluency in the well, usually 13–15 days after transfection, depending on cell recovery after oligo transfection. Fluorescent Green recombinants were distinguished from Yellow parental cells by flow cytometry using narrow bandpass filters (510 ± 15 nm and 540 ± 10 nm) and a differential angular distribution that allowed quantification of each cell without compensation. Recombination frequencies were calculated as the frequency of Green cells among total (Green + Yellow) fluorescent cells (typically ~50,000 cells per trial).

Extended Methods are to be found in the Supporting Information

## ACKNOWLEDGEMENTS

The authors thank Dr. David Knipe for the plasmid pCMV-ICP8, Dr. Jakob Reiser for the plasmid pENTR2B/TREPitt and for advice about lentivirus engineering, Dr. Priya Rai for the 293 T cells and lentiviral plasmids pHR’ CMV 8.2∆R and pCMV-VSV-G, Dr. Adriana Gomez for her help with microscopy, Dr. Kate M. Vignali and Dr. Dario A. A. Vignali for advice with the 2A peptide constructs, and Drs. Guy Howard, Carlos Perez-Stable, and Ramiro Verdun for critically reading the manuscript. This work was supported by the National Institutes of Health [F31-GM089125-01] to MV, funds from the University of Miami Department of Biochemistry and Molecular Biology to RSM, Department of Veterans Affairs, Veterans Health Administration, Office of Research and Development (Biomedical Laboratory Research and Development) Merit Review award [BX000952] to PSC and funds from the Miami VA Geriatric Research, Education, and Clinical Center to PSC and MVC.

## Additional Information

### Author contributions

MV, RSM and PCS designed research; MV and RSM performed research and analysis; and MV, RSM and PCS wrote the paper.

### Disclaimer

The contents do not represent the views of the Department of Veterans Affairs or the United States Government.

